# Spontaneous and Stimulus-Driven Rhythmic Behaviors in ADHD Adults and Controls

**DOI:** 10.1101/2019.12.24.887802

**Authors:** Anat Kliger Amrani, Elana Zion Golumbic

## Abstract

Many aspects of human behavior are inherently rhythmic, requiring production of rhythmic motor actions as well as synchronizing to rhythms in the environment. It is well-established that individuals with ADHD exhibit deficits in temporal estimation and timing functions, which may impact their ability to accurately produce and interact with rhythmic stimuli. In the current study we seek to understand the specific aspects of rhythmic behavior that are implicated in ADHD. We specifically ask whether they are attributed to imprecision in the internal generation of rhythms or to reduced acuity in rhythm perception. We also test key predictions of the Preferred Period Hypothesis, which suggests that both perceptual and motor rhythmic behaviors are biased towards a specific personal ‘default’ tempo. To this end, we tested several aspects of rhythmic behavior and the correspondence between them, including spontaneous motor tempo (SMT), preferred auditory perceptual tempo (PPT) and synchronization-continuations tapping in a broad range of rhythms, from sub-second to supra-second intervals. Moreover, we evaluate the intra-subject consistency of rhythmic preferences, as a means for testing the reality and reliability of personal ‘default-rhythms’. We used a modified operational definition for assessing SMT and PPT, instructing participants to tap or calibrate the rhythms most comfortable for them to count along with, to avoid subjective interpretations of the task.

Our results shed new light on the specific aspect of rhythmic deficits implicated in ADHD adults. We find that individuals with ADHD are primarily challenged in producing and maintaining isochronous self-generated motor rhythms, during both spontaneous and memory-paced tapping. However, they nonetheless exhibit good flexibility for synchronizing to a broad range of external rhythms, suggesting that auditory-motor entrainment for simple rhythms is preserved in ADHD, and that the presence of an external pacer allows overcoming their inherent difficulty in self-generating isochronous motor rhythms. In addition, both groups showed optimal memory-paced tapping for rhythms near their ‘counting-based’ SMT and PPT, which were slightly faster in the ADHD group. This is in line with the predictions of the Preferred Period Hypothesis, indicating that at least for this well-defined rhythmic behavior (i.e., counting), individuals tend to prefer similar time-scales in both motor production and perceptual evaluation.

## 1. Introduction

Rhythm is a central characteristic of many human behaviors. It is expressed through motor actions, such as clapping, walking, dancing and speaking (Bohannon, 1997; McAuley et al., 2006; Pellegrino et al., 2011; Williams & Grant, 1999). Rhythm also benefits perception, since its inherent temporal predictability allows preempting and preparing for upcoming stimuli (Haegens & Zion Golumbic, 2018; Jones, 2019; Nobre & Coull, 2010; ten Oever et al., 2017). Not only is rhythm implicated as important for a wide range of cognitive abilities, but reduced temporal acuity has been linked to difficulties in both attention and language processing (Barkley et al., 1997; Corriveau & Goswami, 2009; Kerns et al., 2001; Noreika et al., 2013; Smith et al., 2002). In particular, individuals with ADHD reportedly display reduced precision on a variety of timing-related tasks and rhythmic behaviors (Dankner et al., 2017; Kerns et al., 2001; Rubia et al., 2003; Smith et al., 2002; Toplak & Tannock, 2005b; Zelaznik et al., 2012). However, to date there is much ambiguity regarding the nature of rhythm-related deficits in ADHD adults and their consequences for daily behavior. The current study is a broad investigation of spontaneous and synchronized motor rhythms and the potential link between them, in adults with ADHD as well as within a typical control population.

In order to elucidate the nature of rhythmic deficits in ADHD, it is helpful to differentiate between two types of rhythmic behaviors: The first is spontaneous production of rhythm, generated internally by the motor system (Fraisse, 1982; Rimoldi, 1951). The second is synchronization to rhythms in the environment, that involves inherent interactions between sensory and motor systems (Repp & Su, 2013). However, the relationship between spontaneous generation of motor rhythms and synchronizing motor actions to external rhythms has not been sufficiently characterized, to date. The current research focuses on understanding the nature of timing deficits in ADHD and whether they can attributed to difficulties in internal generation of rhythms or rather are linked to deficits in sensory-motor interactions.

### 1.1 Internal Rhythmic Preferences

A large body of literature suggests that individuals have a default **Spontaneous Motor Tempo (SMT)**, that is consistently produced during free motion (Fraisse, 1982; Rimoldi, 1951). In humans, spontaneous rhythms are generated in many body parts, including legs, lips, head and hands (Bobin-Bègue et al., 2006; Bohannon, 1997; MacDougall & Moore, 2005; Rose et al., 2020; Todd et al., 2006; Van Dyck et al., 2015) although the SMT literature has predominantly focused on finger-tapping paradigms where participants are instructed to tap their finger “at their most comfortable rate” (Collyer et al., 1994).

Some have argued that these default motor preferences also extend to the realm of auditory perception. Specifically, McAuley and colleagues (2006) report a correspondence between individual SMTs and the tempo that individuals indicate is most ‘comfortable’ for them to listen to, labeled their **Preferred Perceptual Tempo (PPT)**. This finding prompted the *Preferred Period Hypothesis*, suggesting that individuals have a characteristic preferred rhythm, that is generalized across perception and production, and can be attributed to a common internal oscillator (McAuley, 2010; Michaelis et al., 2014; Provasi et al., 2014; Schwartze & Kotz, 2015). Moreover, some have demonstrated facilitation of perceptual sensitivity, and improved precision of motor and inter-personal synchronization at rhythms near one’s SMT (Large & Gray, 2015; Scheurich et al., 2018; Styns & Leman, 2007; Zamm et al., 2015), supporting the notion of endogenous rhythmic preferences. Notably, an individuals’ preferred rhythm seems to change with age, being faster in children and slowing over the years (McAuley et al., 2006). However, to date, attempts to determine the consistency of ‘default’ motor and perceptual preferences within an individual and the degree to which they generalize across rhythmic behaviors have yielded mixed results (Bauer et al., 2015; McPherson et al., 2018; Qi et al., 2019; van der Wel et al., 2009), as these can also be highly affected by situational factors such as time of day, emotional state, physical effort, heart-rate and others (Dosseville et al., 2002; Moussay et al., 2002). Moreover, the manifestation of internal motor and perceptual rhythmic preferences in clinical populations who exhibit timing-related deficits, such as ADHD, has not been studied extensively (Rose et al., 2020). The current study is the first to our knowledge to test the nature of spontaneously generated motor rhythms in ADHD, and their possible link to other perceptual and motor rhythmic behaviors.

### 1.2 Synchronization to External Stimuli

Synchronizing motor actions to sensory rhythms is carried out through action-perception loops, and relies critically on temporal accuracy within both the sensory (Tierney & Kraus, 2013) and motor systems (Schwartze et al., 2011). A potential neural mechanism proposed to underlie sensory-motor synchronization is entrainment of internal neural oscillations, and phase locking between sensory and motor cortices (Haegens & Zion Golumbic, 2018; Morillon et al., 2014; Rimmele et al., 2018; Schroeder & Lakatos, 2009). In contrast to the notion of SMT, synchronization requires the flexibility to generate a broad range of motor rhythms, as dictated by external sources. Therefore, we might ask whether the Preferred Period Hypothesis bears any relevance for synchronization behavior? Some have suggested that, indeed, motor synchronization to external rhythms is facilitated by internal inclinations, which manifests in improved synchronization accuracy for rhythms near one’s preferred rhythm (McAuley et al., 2006; Styns & Leman, 2007). However, in many studies, the range of rhythms to test this was tailored around participants’ personal SMT or was limited to a small number of rhythms. Moreover, individuals may display large variability in their synchronization abilities, irrespective of the specific rhythm tested. Indeed, several studies report findings that are inconsistent with the prediction that synchronization is specifically improved near one’s SMT (McPherson et al., 2018; van der Wel et al., 2009; Van Dyck et al., 2015). Therefore, currently, it is difficult to assert the specific contribution of personal/default rhythmic preferences to motor synchronization capabilities, in explaining individual differences in motor synchronization to external rhythms.

Given the proposed functional role of synchronization in facilitating perception, it is important to understand whether this behavior is indeed biased by internal rhythmic inclinations, as proposed by the Preferred Period Hypothesis. One of the goals of the current experiment was to systematically test this hypothesis, by testing the intra-subject consistency of spontaneous motor and perceptual rhythms, and their relationship to synchronization capabilities across a broad range of tempi, from sub-second intervals (250ms) to suprasecond intervals (2.2 sec). This also enabled us to study which aspects of rhythmic behavior are implicated in ADHD, dissociating between internally-generated rhythms and sensorymotor rhythmic interactions.

### 1.3 Timing in ADHD

Timing-related deficits in ADHD have been demonstrated across multiple timescales and on a variety of tasks including sensorimotor synchronization, duration discrimination and reproduction, verbal time estimation and temporal anticipation (Noreika et al., 2013). Timing impairments have been associated with dysfunctions in neural circuitry known to be important for temporal processing, including the cerebellum, basal ganglia and fronto-parietal networks (Noreika et al., 2013; Valera et al., 2010) as well as neurochemical regulation of dopamine, which is strongly linked to timing functions and motor control (Baldwin et al., 2004; Luman et al., 2015; Rubia et al., 2009). In particular, two main phenomena are consistently observed across many timing-related tasks: A tendency toward hastening, i.e. preferring and producing faster rhythms / shorter intervals, and overall increased intra-subject variability (Barkley et al., 1997; Klein et al., 2006; Meaux & Chelonis, 2003; Rubia et al., 2003; Toplak et al., 2006; Toplak & Tannock, 2005a). However, the majority of these studies all involve forming temporal representations for *external stimuli*, and to date there has been no systematic investigation of spontaneous rhythm production in ADHD.

The goal of the current study was to investigate whether the deficits in rhythmic behavior observed in ADHD are related to imprecision in internal generation of rhythms or whether they should be attributed to reduced acuity in temporal perception and/or sensorymotor interactions? To this end, we conducted a multi-stage experiment in ADHD adults and matched controls, assessing several aspects of rhythmic behavior, including: spontaneous motor tapping (SMT), perceptual preferences (PPT), as well as motor synchronization and memory-based rhythm reproduction. Critically, these tests were performed independently and identically for all participants, whereas in some previous studies evaluation of PPT and motor synchronization-continuation was tailored around participants’ individual SMT. We also emphasize measures of consistency across trials and sessions, and not only center-metrics, as these are central for determining the extent to which motor and perceptual preferences are indeed characteristic and consistent within each individual.

In a previous study, we attempted to replicate reports of a correspondence between SMT and PPT by instructing participants to tap ‘at their most comfortable rate’ and to rate perceived rhythms according to their degree of ‘comfort’ or ‘pleasantness’ (McAuley et al., 2006). However, in that study we failed to find any correlation between motor and perceptual preferences, and participants produced extremely variable results across repeated sessions (see *Data in Brief* report accompanying this paper). We hypothesized that this may be due to the vagueness of the term ‘most comfortable’, which lead participants to interpret it in different ways. Therefore, in the current study we chose to narrow the operational definition of ‘spontaneous rhythms’ and link instructions to a more well-defined internal rhythm – counting (Grondin et al., 1999). We anticipated that giving more specific instructions would produce better correspondence between motor and perceptual rhythms, allowing us to evaluate their manifestation across the two groups tested, while reducing trivial variability related to subjective interpretation of the term ‘comfort’. Additionally, using ‘counting’ instructions would allow us to relate our findings more directly to speech-rhythms, which are often pointed to when considering the ecological relevance of rhythmic preferences and synchronization for everyday behavior (Lagrois et al., 2019; Woodruff Carr et al., 2014).

## 2. Material and Methods

### 2.1 Participants

The experiment included two groups: ADHD and Controls, with 19 participants in each (15 women in each, 4 left handed), aged 21-28 (mean 23 in both). The experiment was approved by the Institutional Review Board of Bar Ilan University. Participants provided written informed consent prior to commencement of the experiment and received compensation for participation.

All participants self-reported normal hearing and no history of neurological disorders (besides ADHD). Participants in the ADHD group presented written diagnosis, from a neurologist or psychiatrist (10 participants) or from certified ADHD clinics. Within the ADHD group, 8 participants regularly took medication on a daily or weekly basis, 3 participants took medication at need and 8 did not take medication at all. However, all participants were instructed not to take medication 24 hours before experiment. Individuals in the ADHD group were not differentiated into distinct subtypes, since in line with the DSM-V guidelines these are no longer considered easily-defined categories (Epstein & Loren, 2013). Administration of an auditory CCPT task confirmed a significant difference between the ADHD and Control groups, complementing their formal diagnosis (see below). Within the ADHD group, 11 participant also self-reported mild to medium-level learning disabilities, which is a common comorbidity with ADHD (Faraone et al., 1998; Luo et al., 2019; Willcutt et al., 2005). The two groups did not differ on years of musical training or sports/dance experience.

### 2.2 Procedures and Stimuli

Participants were seated comfortably in a sound attenuated booth, and heard sounds through headphones (Sennheiser HD 280 pro). Finger taps were recorded using a custom-made tapper based on an electro-optic sensor. All other behavioral responses were collected using a response pad (Cedrus, RB-840). The experiment was programmed and controlled using PsychoPy software (www.psychopy.org). All auditory stimuli were prepared using Audacity and Matlab (Mathworks), and consisted of repetitions of pure tones (440Hz, 30ms with ±5ms ramp up/down), presented at different rates. The experiment consisted of four tasks, performed in interleaved order as shown in Figure 1.

**Figure 1.**
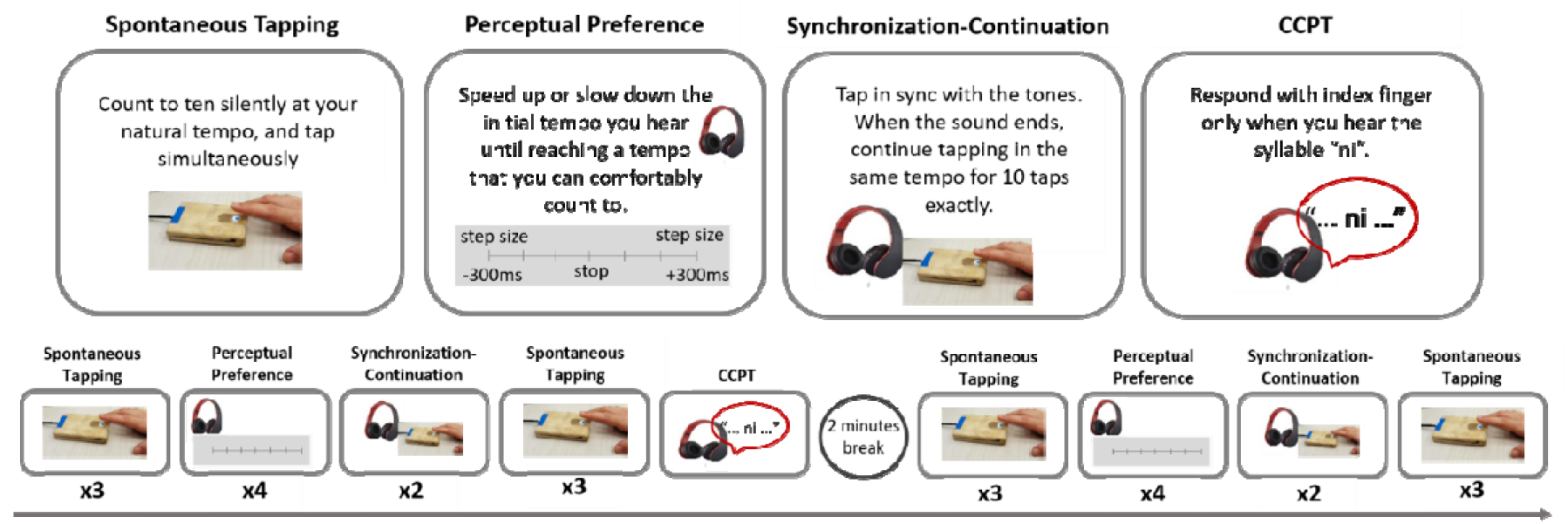
Experimental design. Top: The four tasks performed during the experiment. Bottom: Time line of performing each task and repetition across sessions.

#### Spontaneous Tapping Task

Participants were instructed to tap with the index finger of their dominant hand while counting silently from 1 to 10. After each trial participants received feedback as to the number of taps performed, which ensured that they indeed counted internally. Participants repeated the spontaneous tapping task in four separate sessions throughout the experiment, with each session containing three consecutive tapping trials. These repetitions were used to test for consistencies in spontaneous tapping rates within and across sessions.

#### Perceptual Preference Task

PPT was assessed using a method of limits approach. Participants were presented with sequences of tones and could dynamically adjust their tempo (speed up or slow down) until they reach a tempo that was comfortable for them to count along with. The initial tempo of each trial was either very slow (ISI: 1300 or 1400ms) or very fast (ISI: 250 or 350ms), and participants could change the ISI in discrete intervals of ±30ms, ±100ms or ±300ms. They were instructed to start with using the larger step and then to fine-tune their selection using the smaller steps. Once they had reached a tempo that was comfortable for them to count with from 1 to 5, they pressed a ‘stop’ button. Participants repeated the PPT task in two separate sessions throughout the experiment, with each session containing four consecutive trials (alternating fast and slow initial tempi).

#### Synchronization-Continuation Task

Participants heard a sequence of 10 tones at a particular tempo and tapped along with them (Synchronization stage). Then a stop sign appeared for 1.5 seconds, after which they were instructed to reproduce the tempo they had just heard and continue tapping 10 times (Continuation stage). Participants received feedback after the continuation stage whether they had indeed tapped 10 times, which ensured internal counting. No feedback was given regarding the temporal accuracy of the tapping. Ten different tempi were used, presented in random order (ISIs: 250, 350, 450, 550, 650, 800, 1000, 1400, 1800 and 2200ms). Participants performed the Synchronization-Continuation task in two separate sessions throughout the experiment, and each tempo was repeated twice in each session.

#### Auditory CCPT task

Participants completed an auditory version of the Conjunctive Continuous Performance Test (CCPT), in order complement the formal ADHD diagnosis. We followed the design developed and validated by Shalev and colleagues for Hebrew speaking participants (Shalev et al., 2011). Participants listen to a stream of CV syllables, comprised of all 16 combinations of the consonants (*n, s, b, r*) with the four vowels (*a, e, i, u*), which are clearly distinguishable in Hebrew. Syllables were presented for 200 ms, with ISI ranging 1 second, 1.5, 2 or 2.5 seconds. Participants were required to press the response button with their index finger only when they hear the target syllable */ni/*, which appeared at a frequency of 30%. All non-target items were presented with equal frequency. The test consisted of a 10-minute long block, containing 320 trials preceded by 15 practice trials.

### 2.3 Data Analysis

#### SMT and PPT evaluation

To quantify spontaneous tapping behavior (SMT), we assessed both central tapping measures (mean and median values), as well as consistency measures, within and across trials. All calculations were derived from the Inter-Tap-Intervals (ITIs) in individual trials, and are illustrated in Figure 2. All taps were included in the analysis, without excluding initial taps or extremely long or short ITIs, since we regard any such variability as an integral part of tapping behavior.

**Figure 2.**
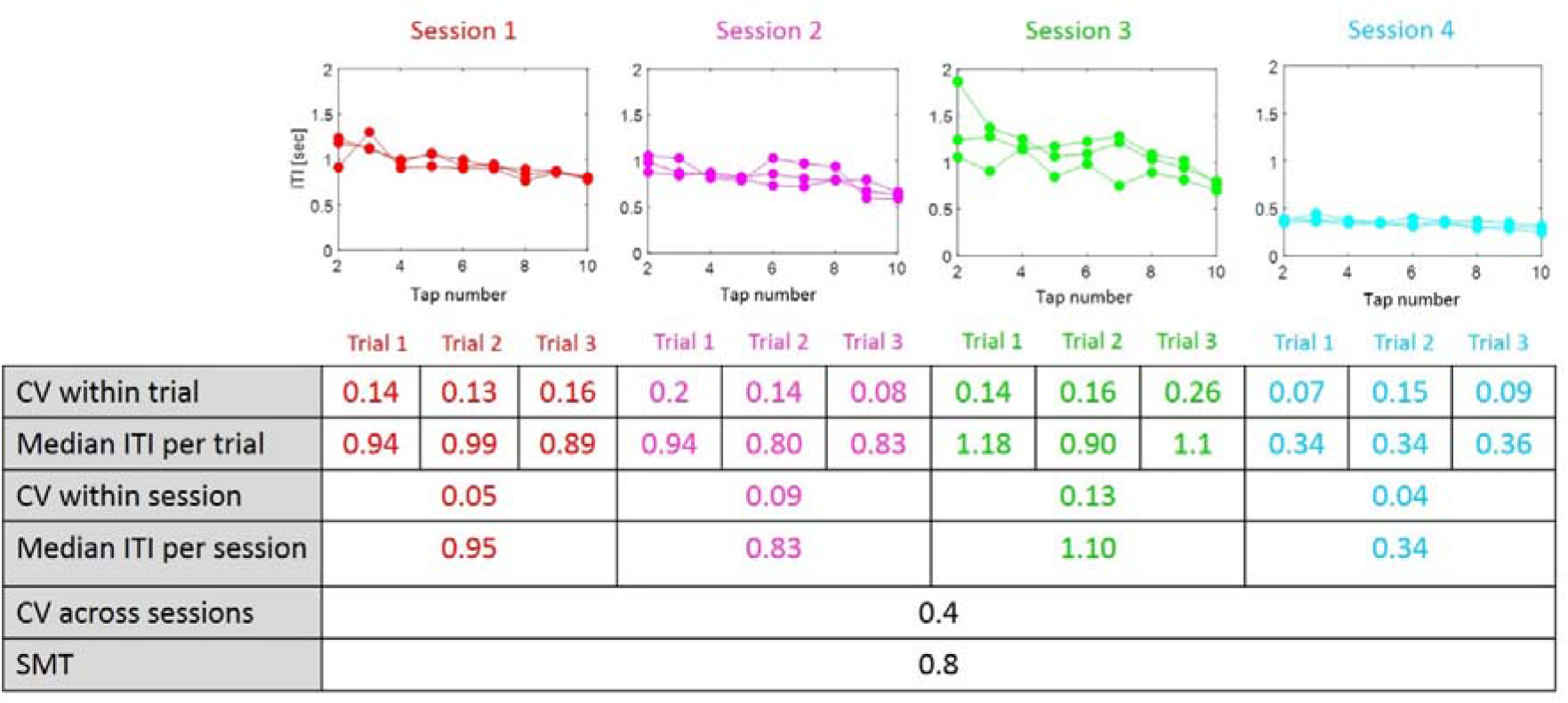
Analysis of Spontaneous Tapping Behavior - example from one participant. *Top:* Tapping ITIs in single trials (10 taps per trial, three trials per session), across all four sessions of the SMT task. *Bottom:* Table summarizing the derivation of central and consistency metrics (median/mean and CV, respectively) from tapping ITIs, within and across session, used for characterizing different aspects of spontaneous tapping.

Consistency measures were all based on the Coefficient of Variation (——, to avoid biases due to differences in tempo and allow comparability across tempi. We specifically calculated the following consistency metrics for each participant:

*Within-trial tapping consistency (CV_within_trial_)*: Represents how isochronous the tapping was within a given trial. This is calculated using the ITIs of all ten taps in a given trial (Figure 2, row 1).
*Within-session tapping consistency (CV_within_session_)*: Represents whether participant replicate the same median rhythm in consecutive trials within a session. This is calculated using the median ITI values from the three trials within each session (Figure 2, rows 2,3):
**Across-session tapping consistency (CV_across_session_)**: Represents whether participant replicate the same median rhythm in different sessions throughout the experiment. This is calculated using the median ITI values from the three sessions (Figure 2, rows 4, 5).

Finally, the average **Spontaneous Motor Tempo (SMT)** was calculated by averaging the median ITIs across all four sessions (Figure 2, row 6).

To asses each individual’s PPT, we averaged all tempi they indicated were ‘comfortable to count with’ across all trials in the Perceptual Preference task (4 trials starting fast; 4 trials starting slow). The consistency of PPT across trials was further calculated as the CV across these 8 tempi.

Differences between the ADHD and Control group in their mean SMT and PPT values and CVs (all types) were tested statistically using Wilcoxon rank sum test (testing for different medians) and Kolmogorov-Smirnov test (testing for different distributions).

#### SMT-PPT correlation

We performed a linear regression analysis to evaluate the correspondence between the SMT and PPT values obtained for each participant, fitting the data to a linear model *y* = *βx* + *β*_0_. The strength of the correspondence was assessed statistically using the r-value associated with the goodness of fit of the model. In addition, we tested whether the slope of the regression line (β) deviated significantly from the diagonal unity line (β=1), by calculating the two-tailed t-test estimation for β, using the statistic t_β_ = (β-1)/SE(β).

Linear regression was performed across all participants, collapsed across groups, and also for each group separately. In addition, to remove potential biases due to outliers, the analysis was also repeated using robust linear regression (Pernet et al., 2013).

#### Synchronization-Continuation

Tapping precision in the Synchronization-Continuation task was evaluated by calculating the ratio between the mean ITI produced in each trial and the prescribed ISI of the stimulus (ITI/ISI) as well as the precision error: **|1 – ITI/ISI|**. We also estimated how isochronous the tapping was by calculating the CV_within_trial_ (similar to the procedure described above for spontaneous tapping). These were calculated separately for the Synchronization and Continuation stages.

For the Synchronization tapping we also calculated the normalized Mean Asynchrony (NMA) which describes the relationship between the timing of each tap (*t_tap_*) relative to the time of the sound itself (*t_sound_*), using the calculation: 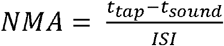 (see Supplementary Material).

We tested if tapping precision-error, tapping isochrony (CV*_within_*) and NMA were modulated by tempo or group using a 10 x 2 mixed-model ANOVA (Within-factor: tempo, Between-factor: Group).

#### Modulation of Synchronization-Continuation performance by SMT/PPT

Last, we tested the prediction of the Preferred Period Hypothesis that tapping in the Synchronization-Continuation task is better near ones’ SMT/PPT, using a nested linear regression analysis aimed at evaluating whether performance at each tempo was modulated by its distance from the participants SMT or PPT (separate analyses). For each participant we estimated two separate regression-lines, for rates either faster or slower than the SMT/PPT, and extracted the slopes (β) estimated for each participant from each of the regressions (see illustration of the procedure in Figure 10). Next, we tested whether the slopes estimated from each regression shared similar signs and if their distribution differed significantly from a null distribution around zero using a sign-test. According to the predictions of the Preferred Period Hypothesis, we would expect to find significantly negative slopes for rhythms faster than ones SMT/PPT, and positive slopes for rhythms slower than ones SMT/PPT. We performed this analysis on the precision-error values in both the Synchronization and Continuation tasks, as well as on the NMA values from the Synchronization task.

## 3. Results

### 3.1 CCPT Results

Before analyzing the difference between groups in the main experiment, we used the CCPT task to verify the distinction between the two groups. Following common practice in the clinical diagnosis of ADHD (Di Martino et al., 2008; Shalev et al., 2011), we quantified the variability in reaction times (RT standard deviation; RT-std) for each participant across the entire task, as a measure for sustained attention. As expected, we found that RT variability was greater in the ADHD group relative to the control group (p=0.033, Wilcoxon rank sum test; Figure 3). This pattern confirmed the ADHD group selection, complementing their clinical diagnoses.

**Figure 3.**
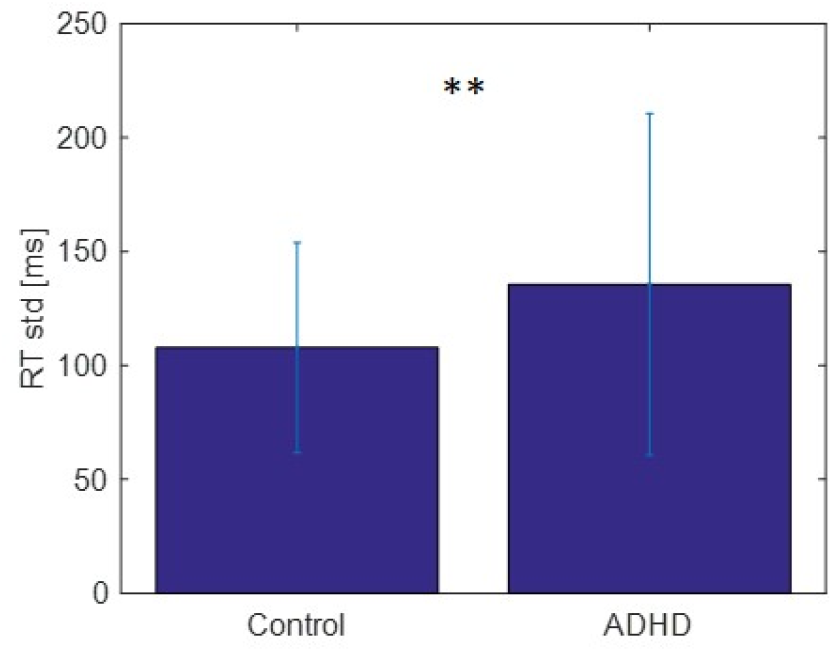
CCPT results. Reaction Time variability (RT std) on the CCPT tasks for the two groups.

### 3.2 Spontaneous Motor Tapping

Both groups exhibited a broad range of median SMT values, between 0.4 - 1.5 seconds ITIs (median = 0.83; Figure 4A). Distribution of SMT values did not differ significantly between the Control and ADHD groups (p=0.24, Wilcoxon rank sum test), although the median SMT tempo in the ADHD group was slightly faster (Figure 4B). SMT distribution was also similar to those obtained in a previous study from our group using the more conventional instructions to “tap at the rate most comfortable to you” (rank-sum test p=0.2; see accompanying Data in Brief report), verifying that the use of a ‘counting’ paradigm here did not substantially alter the range of SMTs produced.

**Figure 4.**
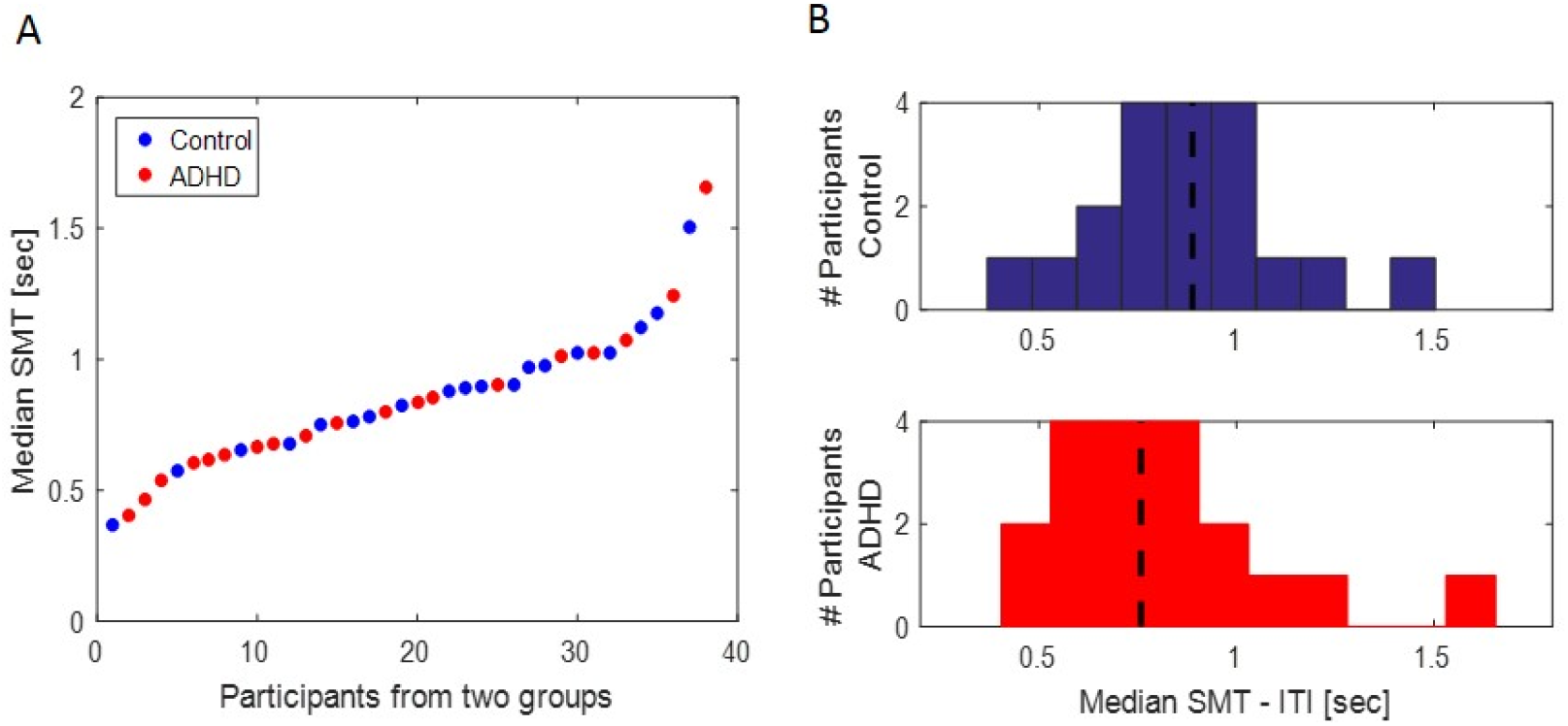
Counting-SMT distribution. A) Counting-SMT values for all participants, color coded by group (blue – control; red – ADHD). B) Distribution of Counting-SMT values by group. Dashed line indicates the median group SMT. No significant differences in SMT distributions was observed between the two groups.

Although previous literature focuses primarily on a single SMT rate per participant, spontaneous tapping within individual participants was in fact often inconsistent, as illustrated in Figure 5C. To characterize this variability, we quantified tapping consistency at three levels: within trial, within session and across sessions.

**Figure 5.**
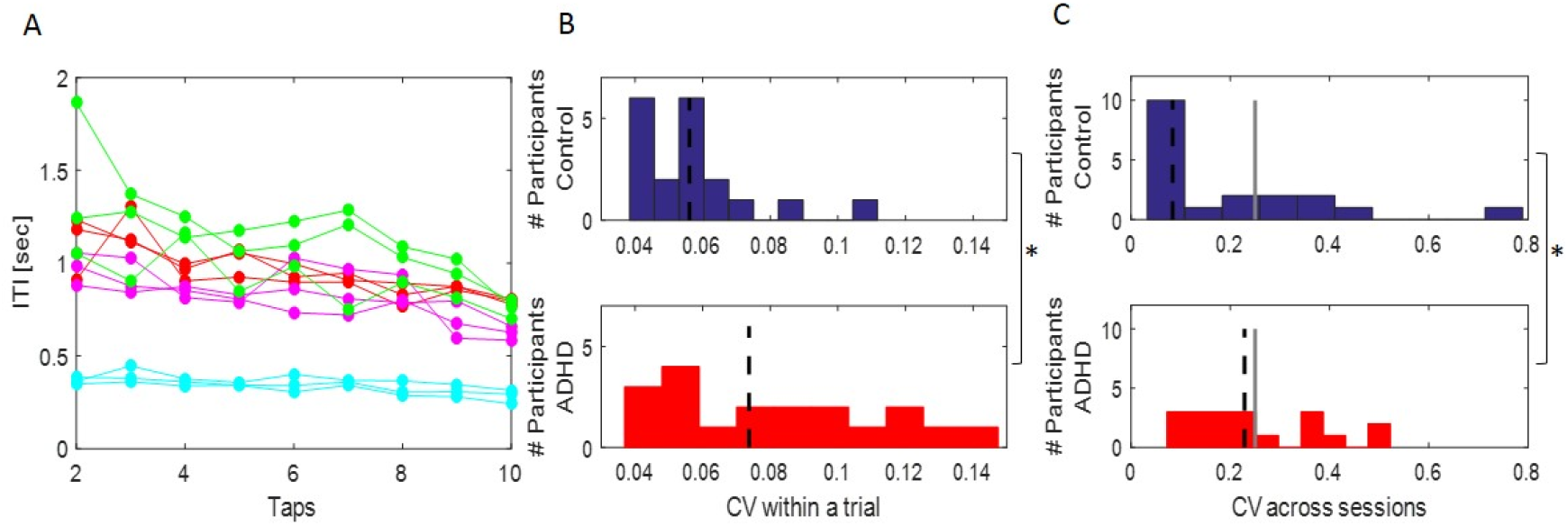
Spontaneous Tapping Consistency. A) Spontaneous tapping times in a single participant across all trials, color coded by session (three trials per session; same data as shown in Figure 2). This example illustrates both within trial variability (e.g., tapping ITIs within the green trials varied within a large range) and across session variability (e.g., tapping in the cyan session was substantially faster than the rest) often observed within participant. B) Distribution of within-trial CV in the Control and ADHD groups. The dashed line represents the groups median. Tapping consistency was reduced in the ADHD group relative to Controls. C) Distribution of across-session CV in both groups. The dashed line represents the groups median, and the gray line indicates the cutoff of CV>0.25 used in a previous study to classify and exclude participants with inconsistent SMT across sessions (McAuley et al. 2006). Inconsistencies in SMT across sessions were significantly more prevalent in the ADHD group relative to controls.

CV_within_trial_ reflects the degree to which spontaneous tapping is isochronous, i.e. whether participants produce a constant rate. We found a significant difference in CV_within_trial_ between the two groups (p<0.03, Wilcoxon rank sum), indicating reduced isochrony in the ADHD group (figure 5B).

The ADHD group also displayed reduced consistency in the median tapping-rates they produced in different trials within the same session (CV_within_session_) and across-sessions (CV_across_session_), as reflected by a more wide-spread distribution (Kolmogorov-Smirnov test; p=0.048 and p=0.018 respectively) and a trend toward higher median CV values (Wilcoxon rank sum test; p=0.054 for both CV_within_session_ and CV_across_session_; Figure 5C). Using a cutoff of CV_across_session_ > 0.25 used previously by McAuley et al. 2006 to exclude participants with inconsistent SMTs, 42% of participants in the ADHD group (8/19) and ~30% of the participants in the control group (6/19) did not produce consistent SMTs across sessions.

### 3.3 Preferred Perceptual Tempo

The tempi that participants indicated as most “comfortable to count with” (mean PPT across sessions) ranged broadly between 0.26-1.46 sec ISI (median 0.84). PPT distribution of the Control group did not differ from those obtained in a previous study from our group using the more paradigm where ratings were given for 10 different rhythms based on how “comfortable” they were (rank-sum test p=1; see accompanying Data in Brief report). This verifies that the changes in the procedure used to evaluate PPTs here, did not substantially alter the range of PPTs. However, comparison between the Control and ADHD groups indicated that median PPT values were significantly faster in the ADHD group (p<0.03, Wilcoxon rank sum test; Figure 6). However, the distribution of PPT rating consistency (CV) did not differ significantly between the groups (Kolmogorov-Smirnov test p = 0.11; Wilcoxon rank sum test, p=0.16; data not shown).

**Figure 6.**
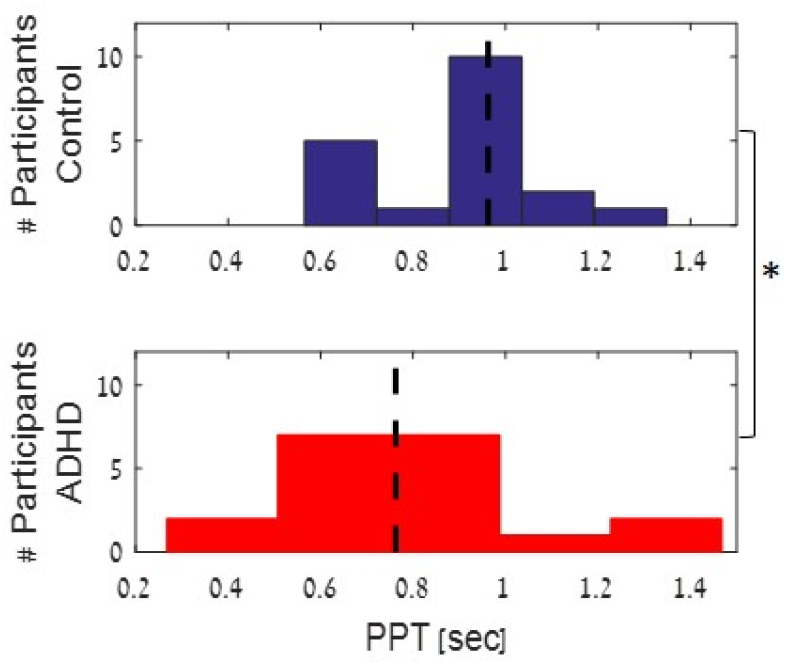
Counting-PPT distribution. Distribution of mean Counting-PPT values across trials in the ADHD and Control groups. The ADHD groups showed overall preference for faster tempi relative to controls.

### 3.4 SMT-PPT correlation

A key claim of the Preferred Period Hypothesis is that auditory perceptual preferences and motor preferences can be attributed to a common underlying oscillator. We tested the relationship between the rhythms generated spontaneously by each participant and the rhythms they indicated as auditory perceptually preferable using a linear regression between the SMT and PPT values (across all sessions). We found that indeed these values were positively correlated (linear regression estimation: y=0.647x+0.288, r=0.678, p<10^−5^, Figure 7A dashed thick line; robust linear regression estimation: y=0.702x+0.27, r=0.73, p<10^−5^, Figure 7A solid line, outliers marked in dashed circles). Despite the positive correlation between the two metrics, the estimated regression line did not fall on the diagonal. Deviation of the estimated linear regression from the unity line was verified statistically by testing whether the estimated slope (β) is significantly different than 1, using a two-tailed t-statistic:

**Figure 7.**
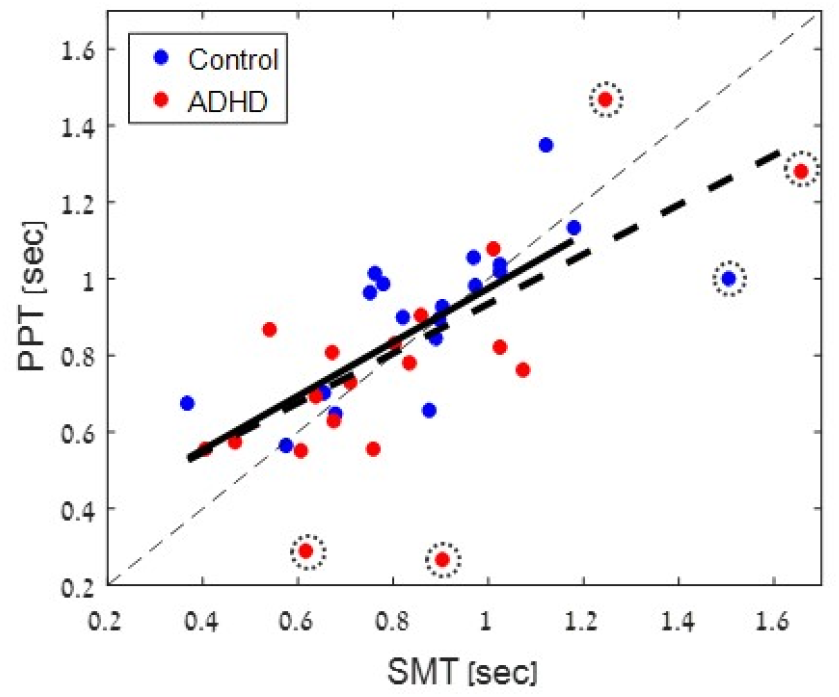
Correspondence between SMT and PPT. Linear Regression between the Counting-SMT and Counting-PPT values within individuals, color coded by group (blue – control; red - ADHD). The dashed thin line is the diagonal unity line. The thick lines indicate the regular (dashed) and robust (solid) linear regression estimation between SMT and PPT values (outliers from robust estimation are circled). Although the two measures are strongly correlated (simple and robust correlation, p<10^−5^), the slope of the regression line is significantly different than 1 (robust p=0.016), indicating a deviation from one-to-one correspondence of preferred motor and perceptual ‘counting’ rhythms.

This yielded a significant effect [tβ(36) = −3.0188, p=0.004; robust correlation: tβ(31) =-2.5389, p=0.016], indicating that we can reject the hypothesis that β=1. Similarly, when fitting the data separately for each group, correlations remained significant yet the regression slope values remained significantly smaller than 1 (control group: y=0.53x+0.44, r=0.67, p<0.002; ADHD group: y=0.67x+0.2, r=0.68, p<0.002).

### 3.5 Synchronization - Continuation

Analysis of performance on the Synchronization-Continuation task showed that Synchronization tapping was substantially more accurate than Continuation tapping, across both groups and all tempi (maximal precision error in Controls: 5% in Synchronization vs. 12% in Continuation; Figure 8). To test for differences in performance across tempi and groups we performed a 2×10 mixed-model ANOVA on the precision error, separately for each task. For Synchronization tapping there was a main effect of tempo [F(9,36)=4.96, p<10^−5^], which was primarily due to slightly increased errors for the two fastest rhythms (250 and 350 ms ISI; Figure 8A), and there was no significant difference between the two groups [F(1,36) < 1], nor was the interaction between Group X Tempo significant [F(9,324) < 1].

**Figure 8.**
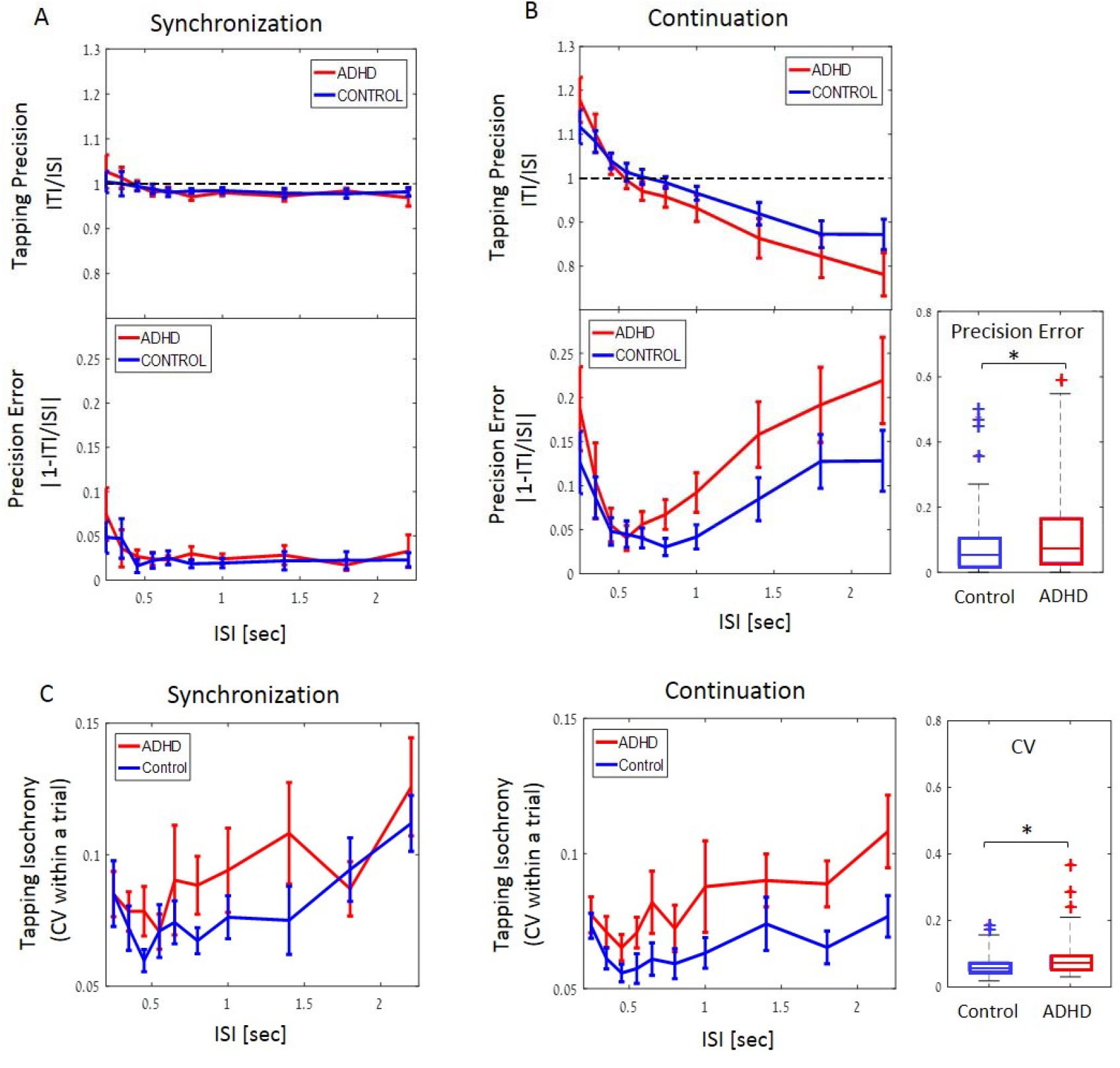
Synchronization and Continuation tapping. A) Tapping precision (top) and absolute precision-error (bottom) for all tempi in the Synchronization task in the Control (blue) and ADHD group (red). B) same as A for Continuation task. Right inset – box plots illustrating the median precision errors and the 25/75^th^ percentiles, averaged across all tempi, for each group. Outliers are indicated by the + sign, (values are considered outliers if they are > 3 times the interquartile range from the top or bottom of the box). C) Within trial tapping variability (CV_within_trial_) in the Synchronization (left) and Continuation (right) across tempi, in both groups, indicating the degree of isochrony. right inset – box plots illustrating the median CV and their distribution, averages across all tempi, for each group. Error bars in all graphs depict SEM.

However, for Continuation tapping a more complex pattern was observed for precision errors, which formed a U-shape [main effect of tempo: F(9,36)=18.04, p<10^−6^; Figure 8B]. In the Control group, optimal performance (mean precision error < 5%) was observed for ISIs between 450-1000ms, and in the ADHD group it was confined to ISIs between 450-650 ms. In the two groups, precision errors increased in opposite directions both faster and slower rhythms, indicating that tapping was slower-than-prescribed for fast rhythms (ISI<450ms), and was faster-than-prescribed for slower rhythms (ISI>1000ms). The main effect of Group was significant [F(1,36)=5.11, p=0.029], indicating that the ADHD group deviated from the prescribed rhythm more than the control group, accentuating the U-shape of the graph, although the interaction between Group X Tempo was not significant [F(9,324)=1.64, p=0.101].

We also assessed whether tapping isochrony (CV_within_trial_) varied as a function of Tempo and Group. There was a main effect for Tempo for both Synchronization and Continuation CV [F(9,36)=4.99; p<0.0002; F(9,36) = 4.53, p<10^−4^, respectively; Figure 8C], with optimal isochrony (least tapping variability) found for rhythms with 350-550 ms ISI, in both tasks. However, here too, a significant effect of Group was found only for Continuation tapping [F(1,36)=4.48, p=0.04], but not in the Synchronization stage [F(1,36) =1.04; p=0.31]. The interaction between Tempo and Group was not significant in either task (F<1.0).

In the Synchronization task we also calculated the Normalized Mean Asynchrony (NMA), describing the relationship between the timing of each tap relative to the time of the sound itself. NMA was found to be negative for all tempi, i.e., the taps preceded the sounds themselves which is consistent with motor-entrainment (as opposed to a sequence of responses to individual sounds). NMA showed a significant modulation across tempi [main effect of Tempo: F(9,36) = 15.4, p<0.0001], but there was no main effect of Group [F(1,36) = 1.51, p=0.22], nor was the interaction between Group x Tempo significant [F(9,324) = 1.37, p=0.19]; see Supplementary Figure S1).

Last, we tested the prediction of the Preferred Period Hypothesis that rhythmic behavior is improved near ones’ individual preferred motor and/or auditory perceptual rhythms. In the Continuation task, this did seem to be the case. Figure 9 illustrates the precision error during Continuation tapping at each tempo for all participants, with individual SMT and PPT values indicated by the cyan and magenta lines, respectively. Precision errors for Continuation tapping at individual SMT and PPT (assessed using linear interpolation between the two nearest tempi) were 2.6% and 2.4% respectively in the Control group, and 5.6% and 4.9% respectively in the ADHD group (Figure 10A). Despite the slightly higher median precision error in the ADHD group, the differences between groups did not reach significance for the SMT (rank sum test, p=0.17), and showed a trend for PPT (rank sum test, p=0.065). When precision error across tempi is aligned relative to each participant’s individual SMT (Figure 10B) or PPT (graph not shown), visual inspection suggests this point serves as a local minima with optimal performance. To quantify whether, indeed, Continuation tapping at these rates is more accurate than at faster or slower rates, for each participant we estimated two linear regressions, describing the relationship between precision error and the distance from ones SMT/PPT, separately for faster and slower rates (see example in Figure 10C). We extracted the estimated slope (β) for each participant from each regression and tested whether their distribution differ significantly from a null distribution around zero (Figure 10D). We found that for rates with an ISI slower than one’s personal SMT/PPT (right portion of the graph), β estimation was significantly positive in both the ADHD and Control groups (sign test, p=0.0192 for SMT and PPT in the control group and p=0.004 and p=0.0007 for SMT and PPT in the ADHD group). For rates with an ISI faster than the SMT/PPT (left portion of the graph), β estimation was significantly negative in Control groups (p=0.004 for SMT and PPT), and in the ADHD group this was true for the PPT (p=0.002), but did not reach significance for SMT (p=0.16). Similar analyses were conducted also for tapping precision error and NMA during the Synchronization task, however these did not show a similar U-shape pattern around participants’ SMT/PPT (see Supplementary Figures S2 & S3).

**Figure 9.**
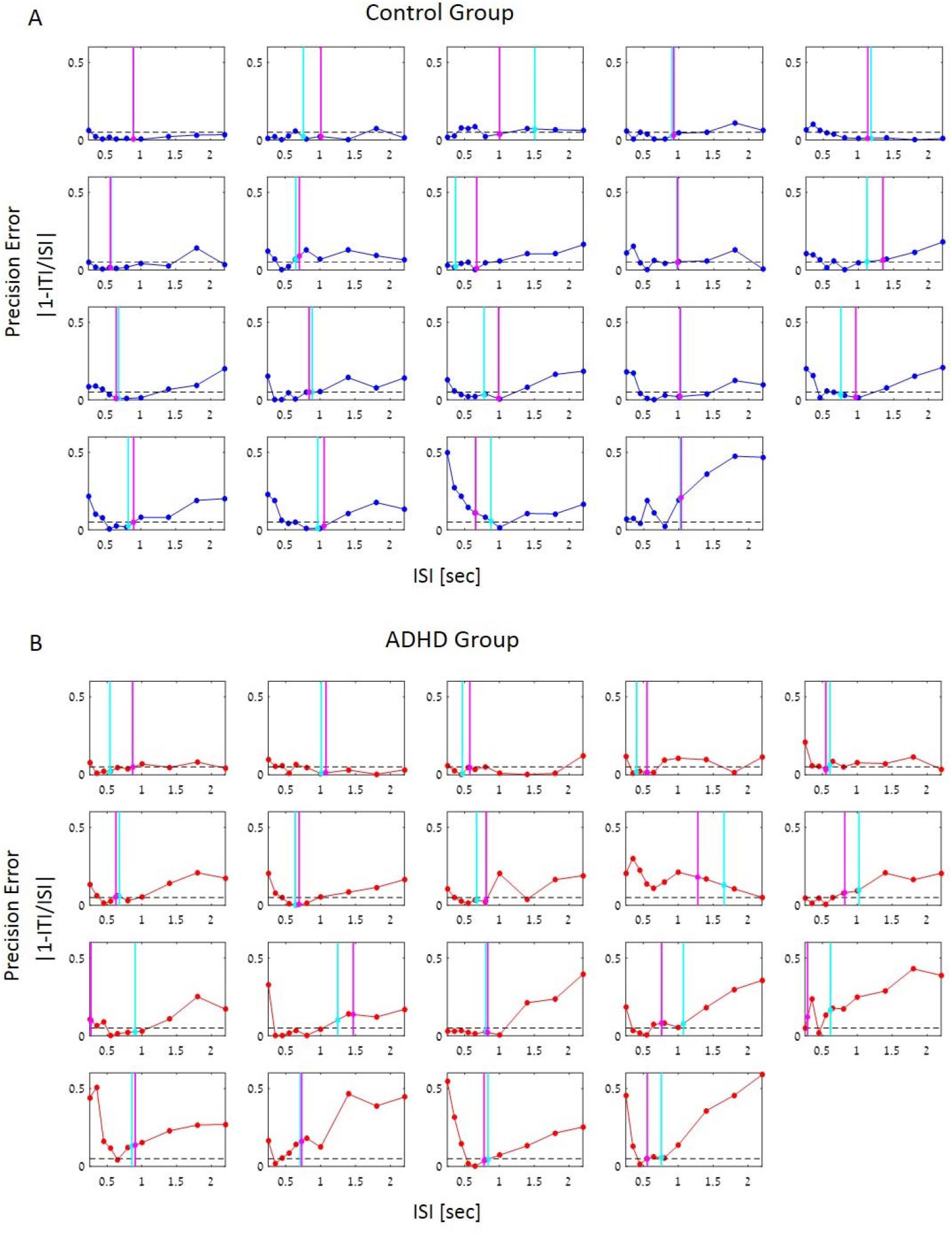
**Precision Error of Continuation tapping for all participants** in the (A) Control and (B) ADHD groups. Subplots are ordered according to the degree of variability in precision errors across tempi. Each participant’s median SMT and PPT are indicated by the cyan and magenta lines, respectively (the precision-error at SMT/PPT was estimated based on linear interpolation of the two nearest tempi, and is marked with a circle of the same color).

**Figure 10.**
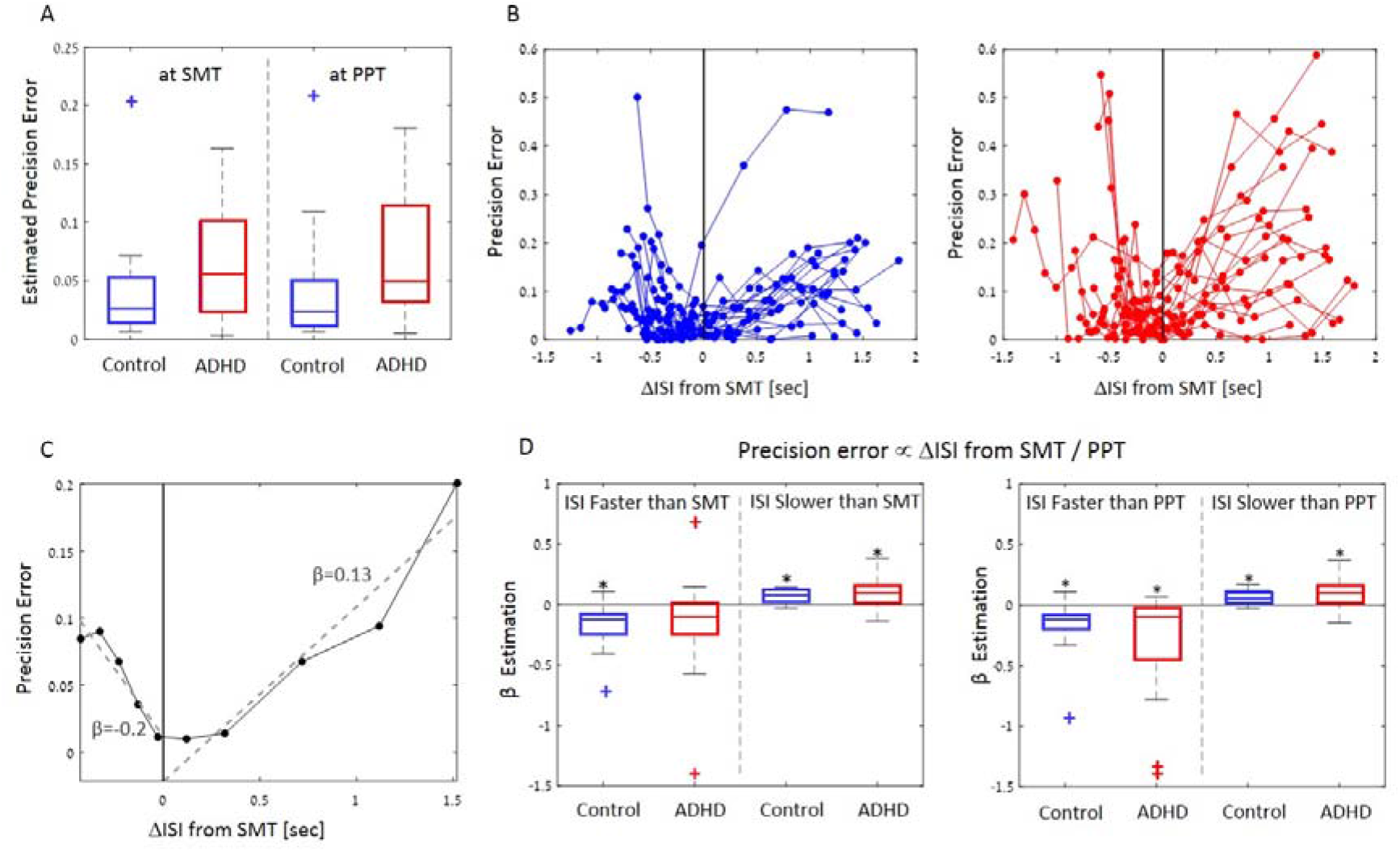
Precision Error of Continuation tapping as a function of distance from SMT/PPT. A) Continuation precision error estimated at each participants’ SMT and PPT, in the Control and ADHD groups. Box plots depict the median of each group and the 25/ 75^th^ percentiles. Outliers are indicated by the + sign (outliers had a value > 1.5 times the interquartile range from the top or bottom of the box). B) Precision error across tempi, aligned to each participant’s individual SMT for the control group (left) and ADHD group (right) C) Example of the linear regression procedure applied to one example participant. A linear fit was performed separately for tempi faster (left) and slower (right) than the participants SMT, and slope values β were extracted for each side. D) Distribution of the estimated β slope values across all participants, showed separately for the analyses conducted relative to the SMT (left) and PPT (right). Consisted with the predictions of the Preferred Period Hypothesis, estimated slopes are negative for rhythms faster than ones SMT/PPT, and positive for rhythms slower than ones SMT/PPT. Box plots depict the median of each group and 25/75^th^ percentiles. Outliers are indicated by the + sign, and defined as in B. Asterisks indicate that estimated β values have a consistent sign across participants (sign test).

## 4. Discussion

The current study provides a broad characterization of spontaneous rhythmic preferences and synchronization to external rhythms in adults with ADHD vs. a typical control population. It sheds light on the nature of rhythmic deficits observed in ADHD, whose primary challenge seems to be with internal generation and representation of rhythms, but are nonetheless accurate at synchronizing to a wide range of external rhythms. It also contributes to a growing body of literature regarding the existence and nature of personal rhythmic ‘defaults’ in motor production and perception. Before discussing the implications of the current findings for understanding nature of rhythmic deficits in ADHD, we first expand on the similarities and differences between the different types of rhythmic behaviors studies here.

### 4.1 Distinctions Between Spontaneous and Synchronized Motor Rhythms

The first aspect of motor rhythms addressed here is the ability to produce isochronous tapping. This was studied under three conditions: Spontaneous finger tapping, paced-Synchronization tapping to an external rhythm, and Continuation tapping at a prescribed rhythm. All three types of behavior manifest in similar motor outcomes, and engage a network of timing-related motor regions (‘temporal hub’), primarily including the supplementary motor area (SMA), inferior prefrontal cortex, the cerebellum, and the basal ganglia (Grahn & Brett, 2007; Matell & Meck, 2004; Merchant et al., 2013, 2015; Schwartze et al., 2011, 2016; Wiener et al., 2010; Witt et al., 2008). However, there are also important distinctions in the processes and neural circuitry they involve.

The isochronous nature of spontaneous, or self-paced, movement has been attributed to the involvement of neural oscillations in motor control (Bichsel et al., 2018; Fraisse, 1982; McAuley, 2010). The tendency toward isochrony is so strong, in fact, that it is quite difficult to produce arrhythmic motor gestures, particularly by the hand, head and legs (Larsson, 2014). In contrast, motor synchronization is not internally generated, but is guided by sensory perception and involves complex interactions between the auditory and motor systems. Even if internal time-keeping is impaired, the auditory input can serve as effective feedback for error correction, allowing listeners synchronize by adjusting their motor tapping to the perceived sounds (Hove et al., 2014; Jacoby & Repp, 2012; Wing & Kristofferson, 1973). Synchronization abilities are usually very good in humans, and can be observed in children as early as 3-4 years old (Fraisse, 1982; Fujii et al., 2014). Synchronization is observed for both simple and complex rhythms in a wide range of tempi (Jacoby & Repp, 2012; Keller & Repp, 2005; Large et al., 2002; Repp & Su, 2013), and the urge to ‘move to the beat’ is often considered near-automatic (Janata et al., 2012; Large, 2000; Nozaradan et al., 2018; Snyder & Krumhansl, 2001; Tal et al., 2017). This is arguably achieved through entrainment of motor neural oscillations to the rhythm of external stimuli to guide and control motor actions (Haegens & Zion Golumbic, 2018; Morillon et al., 2014; Rimmele et al., 2018; Yoles-Frenkel et al., 2016). The idea of entrainment is supported by the well-documented phenomena of negative mean asynchrony in synchronization tasks, observed here as well, which indicates that synchronization is not merely sequential responses to rhythmic input but involves temporal prediction and generalization (Repp & Moseley, 2012).

Thus, the key distinction between spontaneous tapping and synchronization is whether they are guided by an internal or external pace. This is also reflected in the differential neural engagement of components within the ‘temporal hub’ in these two types of behaviors (Bichsel et al., 2018; Chauvigné et al., 2014; De Pretto et al., 2018). Supporting a dissociation between neural mechanisms that are *critical* for self-paced and synchronization tapping, lesions and brain-stimulation studies have shown that the cerebellum and premotor cortex are important for auditory-motor synchronization, but their impairment does not affect generation of self-paced rhythms (Kornysheva & Schubotz, 2011; Schwartze et al., 2016). In addition to motor regions, synchronization tasks also engage bilateral auditory cortex, (Grahn & Rowe 2009, Chen, Zattore and Penhune 2006, Chen, Penhune and Zattore 2008), and difficulties in synchronization are associated with deficits in low-level auditory encoding (Nozaradan et al., 2016; Tierney & Kraus, 2013) and in perception-action coupling (Palmer et al., 2014; Phillips-Silver et al., 2011; Repp & Su, 2013; Schwartze et al., 2016; Sowiński & Dalla Bella, 2013). In contrast, spontaneous self-paced tapping seems to depends more heavily on the basal ganglia, as patients with basal ganglia lesions show higher variability and a lack of isochrony in self-paced tapping, but nonetheless are able to synchronize accurately to a broad range of external rhythms (Schwartze et al., 2011). Intriguingly, clinical populations with basal ganglia lesions and/or Parkinson’s disease suffer from difficulty in maintaining self-generated motion, and yet their movement is greatly improved when hearing rhythms and they are able to synchronize with them quite well (Ghai et al., 2018; Nombela et al., 2013; Schaefer, 2014; Thaut et al., 2001). These findings support a functional and mechanistic dissociation between internally and externally paced motor tapping, and point to the critical role of auditory entrainment in engaging motor oscillations and ‘propelling them into action’, when internal timing cannot be self-maintained.

The case of continuation-tapping is an interesting one, serving as an intermediate between spontaneous and synchronized tapping. Although many studies refer to continuation-tapping as “unpaced” (e.g., Hove et al. 2017), we find that this terminology can be somewhat misleading since continuation tapping is in fact guided by an internal representation or mental imagery of the rhythm previously synchronized to. Some have suggested referring to continuation-tapping as “memory paced”, emphasizing the important role that temporal working-memory plays in perfoming this task, and distinguishing it from spontaneous motor tapping (Chauvigné et al. 2014). Interpretations vary as to whether the brain regions involved in continuation-tapping are more similar to synchronization (Chauvigné et al., 2014) or to spontaneous tapping (Witt et al., 2008). However several studies have pointed specifically to increased activation specifically in the basal ganglia and SMA in continuation vs. synchronization tasks (Koshimori et al., 2019; Lewis et al., 2004; Marvel et al., 2019; Nenadic et al., 2003; Toyomura et al., 2012; Wiener et al., 2010). Given that these regions are also hypothesized to be important for temporal working memory and internal representations of timing (Lustig et al., 2005; Marvel et al., 2019; McNab & Klingberg, 2008; Schmidt et al., 2019; Teki et al., 2017; Wiener et al., 2010), it is possible their engagement in continuation tasks reflects auditory imagery of a rhythm that is used to entrain motor oscillations (Halpern & Zatorre, 1999; Harrison et al., 2018; Schaefer et al., 2014).

### 4.2 Motor Tapping Deficits in ADHD

After establishing the distinctions between spontaneous, synchronized and continuation tapping, we now turn to discuss performance on these tasks in individuals with ADHD. For spontaneous tapping, we found that tapping was more variable and less isochronous in individuals with ADHD relative to controls. To the best of our knowledge only one previous study by Rubia et al. (2003) tested spontaneous tapping in ADHD, focusing on young children (ages 6-12), and similarly found reduced ability to produce isochronous rhythms. Despite their instable spontaneous tapping, sensorimotor synchronization was nonetheless highly accurate in the ADHD group across a wide-range of tempi, in a manner similar to controls. Mean Asynchrony during synchronization was negative for all tempi, indicating true synchronization and entrainment, and here too no significant difference was found between the groups. Previous studies on sensorimotor synchronization to isochronous rhythms in ADHD have yielded mixed results. Results by Tiffin-Richards et al. (2004) are comparable to ours, showing accurate synchronization performance in ADHD across multiple sub-second rates. However, Hove et al. (2017) did find higher synchronization tapping variability in ADHD, at least in two out of four tempi tested (though not adjacent in rate) (see also Pitcher et al. 2002; Ben-Pazi et al. 2003; Rubia et al. 2003; Valera et al. 2010). Therefore, a conservative interpretation would be that variability in synchronization might be slightly enhanced in some cases, but overall synchronization to simple isochronous rhythms is highly accurate in ADHD.

This is in contrast to continuation tapping where the ADHD group tapped less isochronously and their mean tapping rates deviated more substantially from the prescribed rhythm, relative to controls. This is consistent with several previous studies reporting that individuals with ADHD generally exhibit more variability in continuation-tapping vs. synchronization and are also more variable relative to controls (Gilden & Marusich, 2009; Hove et al., 2017; Rubia et al., 2003; Toplak & Tannock, 2005a; Valera et al., 2010; Zelaznik et al., 2012). As discussed above, continuation tapping requires maintaining an internal memory-representation of the rhythm previously synchronized and thus can be considered an intermediate task, sharing some properties with both spontaneous and synchronized tapping.

Thus, the emerging pattern from studying these three types of motor tapping behaviors suggests the following: Individuals with ADHD seem to have an inherent deficit in maintaining internal timing and generating isochronous rhythms based on internal motor-oscillations alone. This manifests both in their reduced ability to produce isochronous tapping in spontaneous and continuation tapping, both of which are internally generated and rely on temporal working memory for maintenance of internal timing representations (Barkley et al., 1997; Mioni et al., 2019). At the same time, auditory-motor entrainment, is preserved in ADHD and is utilized to correct tapping times allowing participant to accurately synchronize to external rhythms, even at extremely slow rates (Ahissar & Assa, 2016; Thaut et al., 2015).

The pattern observed here in the ADHD group, who show impaired self-generated tapping yet optimal performance in the presence of an external pacer, is somewhat analogous to the improvement found in Parkinson’s disease for sensorimotor synchronization (Ghai et al., 2018; Nombela et al., 2013; Schaefer, 2014; Thaut et al., 2001). Although these are highly distinct clinical conditions, they do share some commonalities. Most notably they involve abnormal regulation of dopamine levels (Pine et al., 2010; Vaughan & Foster, 2013) and implicate deficits in basal ganglia function (among other neural substrates) (Curtin et al., 2018; Lu et al., 2019), both of which are important for timing-functions (Allman & Meck, 2012; Pine et al., 2010). As discussed above, spontaneous and continuation tapping share a common reliance on the basal ganglia, which is less critical for synchronization tapping (Chauvigné et al., 2014; Schwartze et al., 2011), and indeed individuals with ADHD show reduced basal-ganglia activation during continuation tapping, coinciding with their reduced performance (Valera et al., 2010). Moreover, a recent PET study showed that synchronized finger tapping to rhythmic sounds modulates dopamine levels in the basal ganglia (Koshimori et al., 2019), suggesting that the behavioral advantages of an external rhythmic pacer are mediated through modulating dopamine-related timing functions. Indeed treatment of ADHD symptoms with Methylphenidate, a dopamine agonist, reportedly improves temporal estimation in individuals with ADHD (Baldwin et al., 2004; Fostick, 2017; Luman et al., 2015; Rubia et al., 2009). Fully exploiting the link between timing deficits in dopamine-related diseases and the role of the basal ganglia, as well as the compensation provided by synchronization to external rhythms is beyond the scope of the current paper. However, these similarities compel further research for understanding the cognitive and neural mechanisms underlying the observed improvement of timing-related functions brought about through synchronization. We note that the current findings that synchronization performance is highly accurate in ADHD relate specifically to simple isochronous rhythms, and may not generalize to more complex temporal tasks such as tapping the beat of complex rhythms or temporal adjustments.

To summarize, our study is the first to directly compare all three types of motor tapping in ADHD adults. Results support a general impairment in the motor production of isochronous rhythms, manifest both in spontaneous tapping and in continuation (reproduction), in a broad range of sub and supra-second rhythms. Nonetheless, we also find that the use of an external isochronous pace-maker as auditory feedback can substantially assist in overcoming the difficulties of internal timing, as the ADHD group successfully adjusted tap-to-tap intervals to maintain the prescribed rate in the synchronization task. The emerging pattern suggests that the temporal deficits in ADHD may be more generally related to difficulties in maintaining an internal representation of a rhythm, or of temporal intervals. Since the current study used a counting-tasks to operationalize rhythmic behavior, these results may also bear significance for processing and producing speech-rhythms (Giraud et al., 2007; Kösem et al., 2018; Poeppel & Assaneo, 2020). Moreover, the current findings contribute to a growing body of literature pointing to benefits of musical training and rhythmic auditory-motor coordination for children with learning disabilities, which is also associated with timing deficits and is highly comorbid with ADHD (François et al., 2013; Gooch et al., 2011; Luo et al., 2019; Thaut et al., 2015; Tiffin-Richards et al., 2004).

### 4.3 Generalizability of Preferred Motor and Auditory Perceptual Rhythms

Another point of interest in the current study was the issue of endogenous rhythmic preferences, as put forth by the *Preferred Period Hypothesis*, postulating that individuals have a characteristic preferred rhythm, generalized across perception and production (Collyer et al., 1994; Fraisse, 1982; McAuley et al., 2006; Michaelis et al., 2014; Schwartze & Kotz, 2015). We tested several key predictions of this hypothesis, primarily the stability of spontaneous preferences over time, the correspondence between auditory perceptual and motor rhythmic preferences, as well as whether spontaneous perceptual and motor preferences transfer to synchronization-continuation tapping (McPherson et al., 2018; Scheurich et al., 2018; Zamm et al., 2015, 2018). Before discussing the results obtained here, it is important to note two important differences in the procedure used here to evaluate individual SMT and PPT relative to most previous studies. First, in some studies, the rhythms used for PPT assessment and for synchronization-continuation testing were tailored specifically around each participants’ personal SMT (e.g., McAuley et al. 2006). This may have biased results toward the central rhythm used, overestimating the correspondence between spontaneous motor and auditory perceptual preferences. To overcome this, in the current study we used a method of limits approach to evaluate individual PPT and tested a broad and constant range of rhythms in the synchronization-continuation task (see also Michaelis et al. 2014). Second, whereas most studies instruct participants to indicate their most ‘comfortable’ rhythm, in the current study instructions were linked to a more well-defined task – counting from 1 to 10 (Grondin et al., 1999). As mentioned above, this choice arguably reduced the within-subject variability observed in our preliminary study where we found no correspondence between SMT and PPT (see accompanying Data in Brief report), which may have been due to the subjective and inconsistent interpretation of what constitutes a ‘comfortable’ rhythm (for similar criticism, see Hinton and Rao 2004). At the same time, it also narrows the interpretations that can be drawn from the current dataset to this more specific operationalization of spontaneous rhythms, referred to in the remaining text as one’s ‘counting rhythm’.

The range of SMTs and PPTs obtained using this more specific ‘counting rhythm’ tasks, was similar to that obtained in previous studies, ranging from ~ 300-1500 ms ISI, with both medians ~800 ms ISI. Moreover, using this approach we also found a strong positive correlation between participants’ median counting-SMT and counting-PPT, in both the ADHD and control groups. It is possible that this correlation was driven, at least in part, by common subvocalization of counting in the two tasks, and thus both may be actually linked to speech-rhythms. However, even so, the correspondence between them was not 1:1, as the regression line deviated significantly from the diagonal unity line. This pattern does not conform to the more extreme prediction put forth by the *Preferred Period Hypothesis*, that default motor and perceptual rhythms should be attributed to a single common oscillator (McAuley et al., 2006; Michaelis et al., 2014). They do support a milder interpretation, that individuals have a default tendency toward either faster or slower rates, at least in the specific context of ‘counting rhythms’. In keeping with previous reports that individuals with ADHD have a tendency toward hastening (Ben-Pazi et al., 2003; Kerns et al., 2001; Meaux & Chelonis, 2003; Rubia et al., 2003), PPTs were faster in this group (although no significant differences in SMTs were observed). However, the current data (and accompanying Data in Brief report) seem to suggest that understanding the nature of individual rhythmic tendencies and their invariance to task and context, still requires additional research.

A second important prediction of the *Preferred Period Hypothesis* is that rhythmic ‘defaults’ be consistent within-subject across repeated testing. Although some studies have looked at within-subject consistency of SMT across trials, this is often limited to a small number of trials/sessions (Schwartze & Kotz, 2015) and individual differences in test-retest reliability are rarely reported. Moreover, a recent study also showed that individuals vary substantially in the degree of isochrony during spontaneous tapping, although in that study SMT replicability across trials/session was not tested (McPherson et al., 2018). Here, we found that ~30% of participants in the control group and ~43% in the ADHD group displayed substantial variability in SMT across sessions. Considering that SMT is known to be affected by situational factors such as time of day, emotional state, physical effort and others (Dosseville et al., 2002; Moussay et al., 2002), these results invite caution in interpreting the SMT measured experimentally through finger tapping as reflecting a singular and unique ‘personal tempo’ (Rose et al., 2020).

A third prediction of the *Preferred Tempo Hypothesis* tested here is whether synchronization-continuation tapping is better at/near one’s preferred tempo. This was achieved testing synchronization-continuation tapping in a broad range of rhythms ranging from sub-second to supra-second rates. We found that, in both groups, synchronization performance was at ceiling for all tempi, and this was not affected by the proximity to one’s preferred range of rhythms. This nicely demonstrates the large range within which individuals (ADHD and controls alike) flexibly modulate motor activity to match the rhythm of external rhythms. Indeed, previous studies have shown similarly good synchronization ability to a wide range of rhythms, on a variety of motor tasks, such as finger tapping (Repp & Su, 2013), dance movements (Phillips-Silver et al., 2011) and running (Van Dyck et al., 2015). Therefore, no additional benefit of personal rhythmic preferences may be required, at least for synchronization to simple isochronous rhythms.

However, this was not the case for continuation tapping. Not only was overall precision lower than synchronization in both groups (London et al., 2019), but results suggests that participants tended to drift with their tapping rate toward a central ‘sweet-spot’ range of rhythms, where continuation tapping was optimal, producing a U-shape for precision-error across tempi (Allman et al., 2014). At the group-level, this ‘sweet-spot’ ranged broadly between 450-1000ms ISI, a range that includes the group-median SMT/PPT values. Moreover, when analysing continuation-tapping performance relative to an individual’s median SMT/PPT (acoss sessions), we found larger percision errors at rates both faster and slower than one’s SMT/PPT. In other words, at least for most individuals the SMT/PPT seem to fall within the ‘sweet spot’ for optimal continuation tapping, and when attempting to perform memory-based tapping for faster/slower rhythms, de-facto tapping rates drifted towards producing rhythms in the vicinity of one’s SMT/PPT. This pattern is consistent with the notion that individuals have a preferred range of rhythms, within which complex rhythmic behavior and perceptual sensitivity is optimal (McPherson et al., 2018; Phillips-Silver et al., 2011; Scheurich et al., 2018; Schurger et al., 2017; Zamm et al., 2015, 2018), and extends these findings to memory-based continuation tapping. Interestingly, similar U-shape patterns were observed for the ADHD and control groups around their SMT/PPTs, eventhough the SMT/PPT rhythms themselves were slightly faster in the ADHD groups.

Taken together, the current study offers several important nuances regrading the empirical validation of the Preferred Period Hypothesis. Supporting the Preferred Period Hypothesis, when looking at the median value of spontaenously-generated motor and perceptual ‘counting rhythms’, across multiple trials and sessions, we find that they are broadly correlated and that memory-based continuation tapping is optimal in their vacinity. At the same time, we also observe substantial within-subject test-retest variability in the spontaneous rhythms produced across sessions, which is often overlooked in similar studies. Moreover, synchronization accuracy was maintained regardless of individual SMT/PPT, and at least in some individuals this was true for continuation accuracy as well. Therefore, rather than asserting that individuals have one preferred rhythm that underlies and guides rhythmic perception and production, we propose adopting a more flexible perspective: individuals may have a general tendency toward a particular range of rhythms, yet de-facto behavior can also be highly flexible, allowing the production of a broad repertoire of rhythms with high accuracy.

## 5. Conclusions

This study highlights two important points regarding rhythmic motor tapping behavior, in ADHD adults and controls. *First*, our findings support a mechanistic distinction between self-generated and synchronized rhythmic tapping. Synchronized tapping is highly accurate and flexible across a large range of sub-second to supra-second rhythms, in both the control and ADHD population. However, maintaining an isochronous rhythm without an external pacer, guided either by temporal working memory or internal counting, is substantially more difficult, particularly for individuals in the ADHD group. These results shed new light on the specific type of temporal processing that are implicated in ADHD adults, and indicate that the use of external pace-makers can assist in overcoming some of these difficulties. *Second*, our results support the existence of preferred time-scales for both perception and production, at least in the specific context of ‘counting rhythms’ tested here, although there is also substantial trial-to-trial and cross-modality variability. At the same time, individuals are not limited by their ‘default’ tendencies, as auditory-motor synchronization is highly adaptable across a wide range of time scales. This flexibility is arguably extremely important for many real-world aspects of behavior, including attention, communication skills and social interactions (Haegens & Zion Golumbic, 2018; Jones, 2019; Lagrois et al., 2019; Nobre & Coull, 2010; Tierney & Kraus, 2014). Further research is necessary to fully understand the relationship between internal rhythmic preferences and the ability to adapt behavior to the wide range of complex rhythms encountered in more ecological contexts.

## Acknowledgements

This work was supported by FP7-CIG grant (2013-631265), and the ISF I-Core Center for Excellence 51/11. We would like to thank Dr. Lilach Mevorach for helpful consultations regarding timing deficits in ADHD, and procedures for ADHD recruitment.

## Supplementary Material

**Figure S1.**
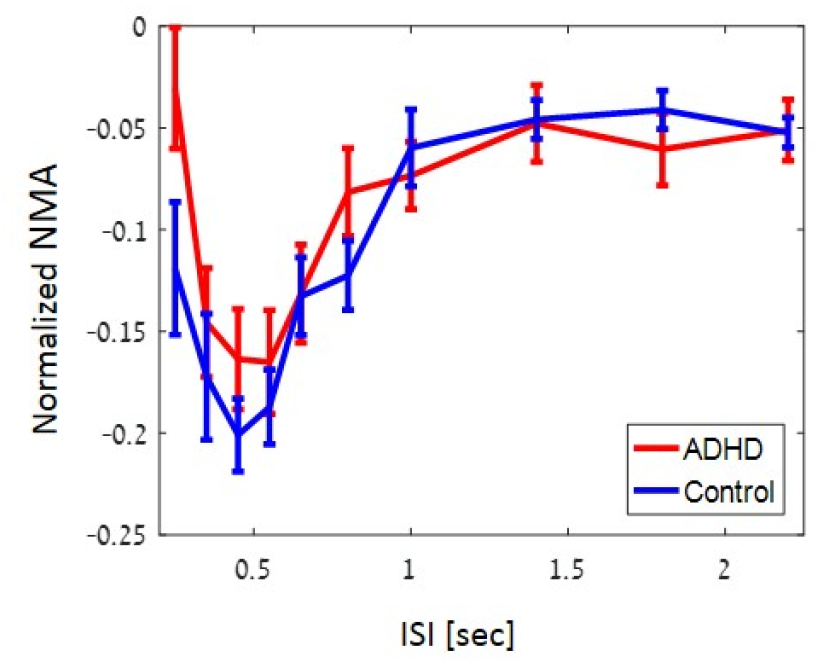
Negative Mean Asynchrony (NMA). Normalized NMA was negative for all tempi, indicating that the motor taps preceded the sounds themselves. NMA size was significantly modulated by Tempo [F(9,36) = 15.4, p<0.0001], with larger NMA values for rates between 350-550ms ISI, and NMA reaching a relatively stable value of −5% for supra-second rates. No differences were found between the groups, nor was the interaction significant.

**Figure S2.**
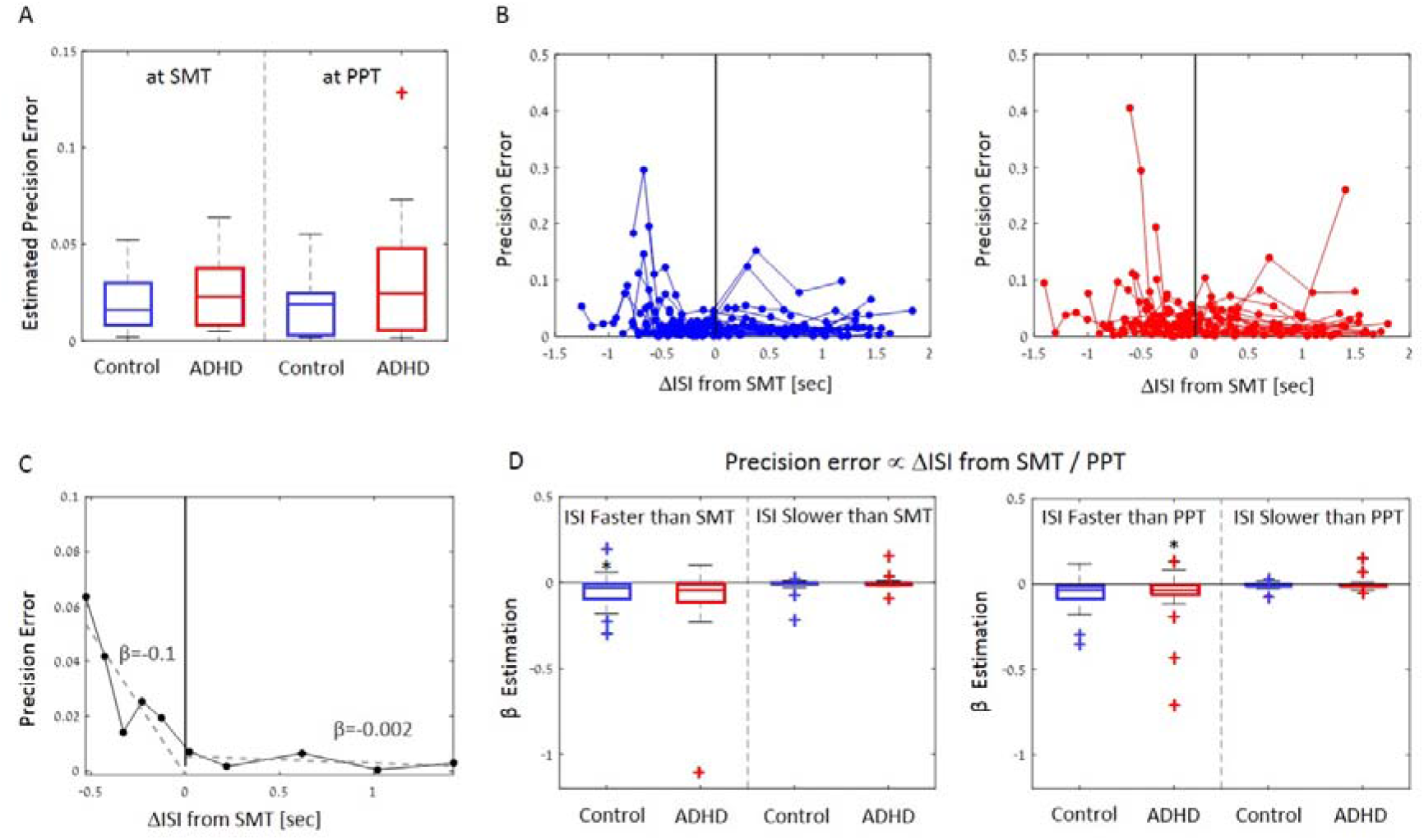
Precision Error of Synchronization tapping as a function of distance from SMT/PPT. A) Synchronization precision error estimated at each participants’ SMT and PPT, in the Control and ADHD groups. Box plots depict the median of each group and the 25/ 75^th^ percentiles. Outliers are indicated by the + sign. B) Precision error across tempi, aligned to each participant’s individual SMT for the control group (left) and ADHD group (right) C) Example of the linear regression procedure applied to one example participant. A linear fit was performed separately for tempi faster (left) and slower (right) than the participants SMT, and slope values β were extracted for each side. D) Distribution of the estimated β slope values across all participants, showed separately for the analyses conducted relative to the SMT (left) and PPT (right). Box plots depict the median of each group and 25/75^th^ percentiles. Outliers are indicated by the + sign. Asterisks indicate that estimated β values have a consistent sign across participants (sign test). Rhythms faster than the SMT/PPT slopes tended to be consistently negative (for SMT: sign test p=0.01 and p=0.06 for Controls and ADHD, respectively; for PPT: sign test p=0.06 and p=0.01 for Controls and ADHD, respectively). However, rhythms slower than the SMT/PPT were not modulated consistently as a function of their distance from SMT/PPT. This pattern suggests that the SMT/PPT does not necessarily constitute a ‘sweet spot’ for Synchronization tapping, but rather that synchronization becomes increasingly inaccurate as rhythms get faster.

**Figure S3.**
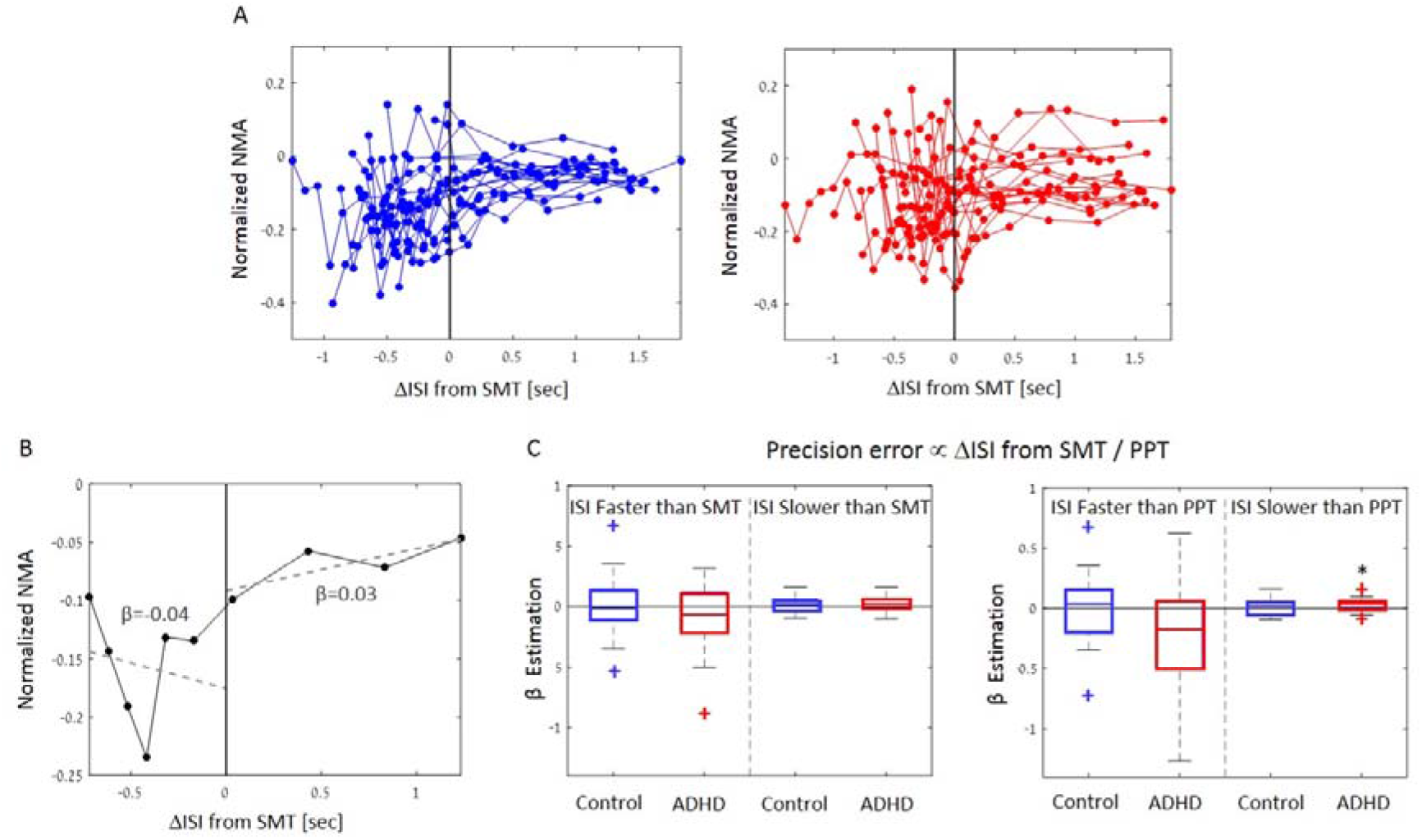
Normalized NMA of Synchronization tapping as a function of distance from SMT/PPT. A) Normalized NMA across tempi aligned to each participant’s individual SMT for the control group (left) and ADHD group (right) B) Example of the linear regression procedure applied to one example participant. A linear fit was performed separately for tempi faster (left) and slower (right) than the participants SMT, and slope values β were extracted for each side. C) Distribution of the estimated β slope values across all participants, showed separately for the analyses conducted relative to the SMT (left) and PPT (right). Box plots depict the median of each group and 25/75^th^ percentiles. Outliers are indicated by the + sign. Only one comparison showed consistent modulation of NMA as a function of the distance from SMT/PPT (rhythms slower than PPT in the ADHD groups, sign test p=0.019). Despite the observed variation in NMA across tempi (Figure S1), this analysis suggests that NMA is not consistently affected by the proximity of a particular rhythm to an individual’s SMT/PPT.

